# Fertility Preservation of Vacuum-Dried Ram Spermatozoa Stored for Four Years at Room Temperature

**DOI:** 10.1101/2024.10.31.621253

**Authors:** Margherita Moncada, Martina Lo Sterzo, Luca Palazzese, Francesca Boffa, Debora Agata Anzalone, Domenico Iuso, Marta Czernik, Luisa Gioia, Federica Turri, Beatrice Mislei, Diego Bucci, Jacques Bonnet, Marthe Colotte, Sophie Tuffet, Michał Bochenek, Pasqualino Loi

**Affiliations:** Department of Veterinary Medicine, University of Teramo, 64100 Teramo (TE), Italy; Institute of Genetics and Animal Biotechnology of the Polish Academy of Sciences, Jastrzebiec, 05-552, Warsaw, Poland; Millio s.r.l. Innovative Start-up of the University of Teramo, 64100 Teramo (TE), Italy; Department of Bioscience and Technology for Food, Agriculture and Environment, University of Teramo, 64100 Teramo (TE), Italy; Institute of Agricultural Biology and Biotechnology (IBBA), National Research Council (CNR), 26900 Lodi (MI), Italy; AUB-INFA National Institute of Artificial Insemination, University of Bologna, 40057 Bologna (BO) Italy; DIMEVET, University of Bologna, 40064 Bologna (BO), Italy; Laboratoire de Recherche et développement, Imagene Company, 33600 Pessac, France; Université de Bordeaux, Institut Bergonié, INSERM, 33076 Bordeaux Cedex, France; Imagene, plateforme de production, Genopole, 91000 Evry, France; Malopolska Centre of Biotechnology, Jagiellonian University, 30-387 Krakow, Poland

**Keywords:** Assisted reproductive technology, ruminant, biobanking, *in-vitro*, anhydrobiosis

## Abstract

Dry storage at room temperature (RT) could simplify spermatozoa banking. Here, we explored DNA stability and *in vitro* and *in vivo* development of embryos derived from vacuum-dried encapsulated (VDE) ram spermatozoa stored for four years or after accelerated aging. While some genomic damage was detected at time 0, DNA fragmentation increased from 3.32±3% (time 0) to 37.64±4% (4 years). A decrease in blastocyst rate was observed after four years of storage and 6.7 years of simulated storage (10.2% and 9% *versus* 13.16% at time 0). Embryo quality, assessed based on *Cdx2* and Inf-τ gene expression, declined over time. Only two of the 23 embryos transferred into synchronized ewes were implanted but were lost by day 40.

In conclusion, dry spermatozoa generated blastocysts after four years of RT storage, but their post-implantation development was impaired. Optimization of the water extraction and storage conditions could better preserve the spermatozoa’s DNA integrity, resulting in improved embryo quality, compatible with development to term.

## INTRODUCTION

In assisted reproductive technologies (ART), spermatozoa from most mammalian species, including humans, are cryopreserved in liquid nitrogen (LN) before being used for artificial insemination (AI), *in vitro* fertilization (IVF) or intracytoplasmic sperm injection (ICSI). However, cryostorage comes with great resource consumption, economic costs, and environmental impact due to its high carbon footprint [1]. Therefore, alternative methods, such as dry storage, were suggested as environmentally friendly and cost-effective alternatives for banking sperm for various purposes, including biodiversity preservation [2].

Indeed, compared to other cell types, spermatozoa exhibit lower water content and a unique DNA packaging. Therefore, it is not surprising that they have been the first cell type to be successfully dried [3]. Even if lyophilized spermatozoa do not retain motility due to membrane damage, their fertilization capacity can be assessed by ICSI. In fact, lyophilized mouse spermatozoa led to embryonic development after injection into mature oocytes, resulting in the birth of the first viable pups [3]. This breakthrough was later replicated in other species, confirming the possibility of obtaining healthy offspring from lyophilized spermatozoa in rabbits [4], rats [5], horses [6], and hamsters [7]. For other large mammals, like pigs [8], cattle [9], and sheep [10], lyophilized spermatozoa were shown to contribute to *in vitro* embryo development, but healthy offspring production is still lacking.

One potential reason for this limitation could be the differences in sperm nuclear organization among species. It has been demonstrated that some domains of the spermatozoa’s DNA retain a nucleosomal organization, likely making them highly vulnerable to desiccation stress. The proportion of nucleosomal-arranged DNA varies among species; mouse spermatozoa retain only 1% of nucleosomal organization [11], humans and cattle spermatozoa retain around 10–15% [11], and nothing is known about other species. Therefore, nuclear organization in mouse spermatozoa explains the success of obtaining a high rate of live offspring when using dry spermatozoa; conversely, achieving comparable results in farm animal species, like sheep, is still out of reach.

The second issue that naturally follows when considering dry spermatozoa biobanking is the storage conditions. Lyophilized spermatozoa are often stored at low temperatures (–20 to –80°C) [12]. However, room temperature (RT) storage would be the most advantageous option for dry genetic banks. Notably, there is limited research on RT preservation, with contributions from murine species [13,14]. Given the greater similarity in DNA compaction between these species and humans, it would be very interesting to have the same information regarding large mammals.

After an exhaustive literature review, we identified vacuum drying followed by encapsulation (VDE) as a method for long-term preservation of nucleic acids in the anhydrous state at RT [15]. Our recent study on applying VDE to ram spermatozoa revealed a preservation of fertilization capacity comparable to that of conventional freeze-drying. This effect was observed in spermatozoa preserved for two years at RT [16]. Spin-drying offers significant advantages over freeze-drying; it is faster (3 h vs. 24 h), more accessible, cheaper, and without the need for liquid nitrogen. Due to these advantages, we opted to further explore spin-drying in our research. Despite its potential, reports on successful offspring production using vacuum-dried spermatozoa are scarce in the literature. To address this gap, we conducted this original research, aiming to contribute valuable insights to reproductive science and potentially advance the use of vacuum-dried spermatozoa in ART.

To this extent, this work explored the DNA stability, fertilization capability, and *in vitro* and *in vivo* development of embryos derived from vacuum-dried ram spermatozoa stored for four years at RT. Furthermore, we thermally stressed dry ram spermatozoa following a method previously applied to mice spermatozoa to mimic a longer storage period and assess spermatozoa’s protein, membrane, and DNA damage over time [12].

Analytical endpoints included the assessment of the developmental rate of *in vitro* matured oocytes fertilized by ICSI with dry spermatozoa, DNA damage (SCSA), and the expression of major pluripotency genes. Selected blastocyst-stage embryos were transferred into previously synchronized recipient ewes for development to term. Our results showed that while the drying procedure was compatible with sperm genomic stability, the same could not be said for long-term storage, which was associated with DNA damage. We believe that this DNA damage was responsible for the poor quality of the embryos, particularly for trophoblastic-specific gene expression, and the developmental failure following embryo transfer. Our work is the first to explore the pre- and post-implantation development of embryos derived from dry spermatozoa—stored for a considerable time at RT—in a large animal, the sheep. We are confident that further technical developments will bring the technology closer to practical application, i.e., biobanking for biodiversity preservation, as well as to assisted reproduction in humans.

## RESULTS

### Impact of VDE on Sperm Viability and DNA Integrity

The membrane integrity (propidium iodide [PI]) results showed that all spermatozoa were dead after VDE (Figure 1b). Pisum sativum agglutinin (PSA) staining categorized the spermatozoa into two: those with intact (IA) or damaged (DA) acrosome. The ratio between sperm with DA and total sperm number was evaluated to assess the effects of the VDE technique on sperm structure (Figure 1d). The analysis revealed the following percentages of sperm with damaged acrosomes: VDE 0, 81.77±1.6%; VDE 4y, 70.17±2.4%; 100C, 87.96±1% (Figure 1d). This observation suggests that the VDE technique does not maintain sperm integrity and that their vitality is completely compromised. Furthermore, the data show that the duration of RT storage did not lead to a significant and evident increase in sperm membrane damage and acrosome loss (Pearson index=0.7) (*P*>0.005) (Figure 1d). The effectiveness of the VDE technique on DNA integrity at time 0 and after RT preservation for five months and four years has been evaluated by SCSA analysis. The results showed an increase in the DNA fragmentation index (DFI%) over time (VDE 0, 3.32±3%; VDE 5m, 22.91±19.45%; VDE 4y, 37.64±4%) (Pearson index= 0.88, P>0.05). For comparison, the DFI% of frozen spermatozoa (FS) was 3.84±1%. Comparisons among treatments found significant differences (FS vs. 4y, *P*=0.01; VDE 0 vs. VDE 4y, *P*=0.002; Table 1, Figure 1e).

**Table 1.** SCSA outcomes: DNA fragmentation at various time points during RT storage.

| Variable | Experimental groups |  |  |  |  |  |
| --- | --- | --- | --- | --- | --- | --- |
|  | FS | VDE 0 | VDE 5m | VDE 4y | 100C | EG effect |
| DFI% | 3.84±1<sup>a</sup> | 3.32±3<sup>b</sup> | 22.91±19.45 | 37.64±4<sup>ab</sup> | 56.91±43.21 | *P*<0.05 |

Values are presented as least square means and standard error of the means. The same superscript letters indicate significant differences between experimental groups at *P*<0.05
EG, experimental group. %DFI, DNA fragmentation index. RT, Room Temperature. FS, frozen semen; VDE 0, VDE spermatozoa at time zero; VDE 5m, VDE spermatozoa stored at room temperature (RT) for five months; VDE 4y, VDE spermatozoa stored at RT for four years; 100C, VDE spermatozoa stored at RT for four years and then incubated at 100°C for ten minutes.

**Figure 1.**
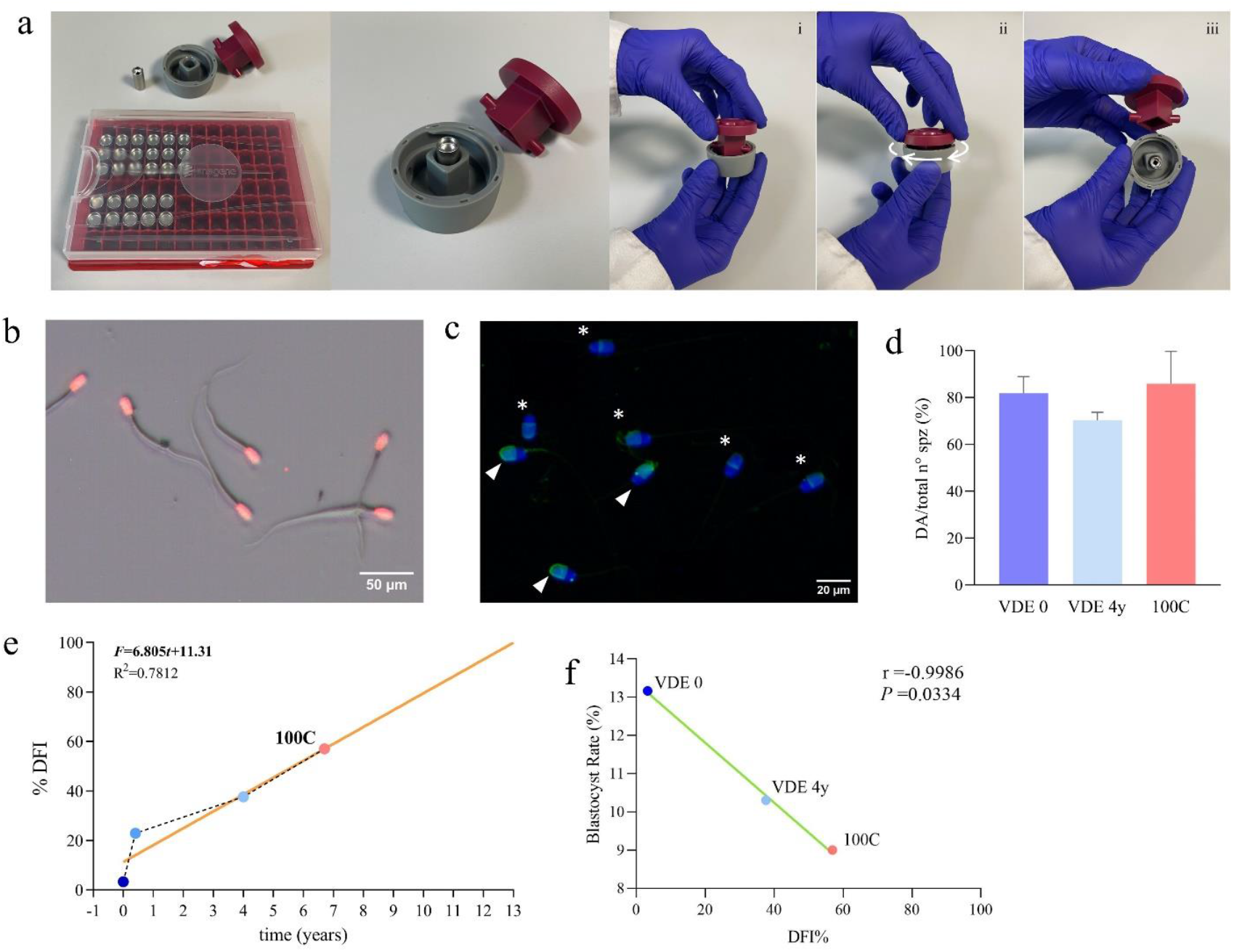
Impact of VDE on Sperm Viability and DNA Integrity. **a**. The two left images show the VDE minicapsules and the ad hoc shellOpener; the three right images show, from left to right, the sequence of the minicapsules opening process; **b**. PI-stained rehydrated VDE spermatozoa. Red staining indicates that all spermatozoa were with damaged membranes; **c**. Rehydrated VDE spermatozoa stained with PSA (green) and DAPI (blue). Triangles indicate intact acrosome (IA) spermatozoa, and asterisks indicate damaged acrosome (DA) spermatozoa; **d**. The ratio of spermatozoa with damaged acrosome to the total number of spermatozoa; **e**. DNA fragmentation index of VDE 0, VDE 5m, and VDE 4y, plotted to construct a linear regression curve of the change in DNA fragmentation over time; the orange line indicates the linear regression decay curve (*F* = 6.805*t* + 11.31, R^2^ = 0.7812); 100C, plotted on the curve, indicates that its DFI% corresponds to storage for approximately 6.7 years at RT; **f**. A graph showing the negative correlation between the blastocyst rate and spermatozoa DFI% at VDE 0, VDE 4y, and 100C (r = –0.9986, *P* = 0.0334). VDE 0, vacuum dry encapsulated (VDE) spermatozoa at time 0; VDE 4y, VDE spermatozoa stored for four years at room temperature (RT); 100C, VDE spermatozoa stored for four years at RT and then incubated at 100°C for 10 minutes; *F*, DFI%; *t*, time; r, Pearson’s coefficient.

The DNA fragmentation data for the VDE 0, VDE 5m, and VDE 4y groups were used to create a curve to visualize the DNA degradation over time. The linear regression equation was *F= 6*.*805t + 11*.*31* (where F = DFI% and t = time) and *R*^*2*^ *= 0*.*7812*. Extrapolation based on this linear curve suggested that after approximately 13 years of storage at RT, 100% of the spermatozoa would be damaged.

Furthermore, the DFI% after exposing the sperm to thermal stress at 100°C for ten minutes (56.91±43.21%) was plotted on the same straight line, indicating that such DNA damage would be reached after 6.7 years of storage (Table 1 and Figure 1e).

Statistical analysis indicated a negative correlation between the DFI% at VDE 0, VDE 4y, and 100C and the corresponding blastocyst rates (Pearson index = –0.9986, *P*<0.05; Figure 1f).

### ICSI Outcomes Remained Stable Over Four Years of Storage at RT

Samples stored at RT for 0, 2, 3, and 4 years were used for ICSI to determine whether VDE spermatozoa retain their fertilizing ability. FS semen was used as a control. The results are shown in Table 2.

**Table 2.** Embryo development of sheep oocytes injected with FS or VDE spermatozoa stored at RT for 0, 2, 3, or 4 years and after 100C treatment.

| Group | No. oocytes | 2-Cell (%) | Blastocysts (%) |
| --- | --- | --- | --- |
| FS | 29 | 8/29 (27.59)<sup>a</sup> | 8/29 (27.59)<sup>a</sup> |
| VDE 0 | 38 | 14/38 (36.85) | 5/38 (13.16) |
| VDE 2y | 53 | 25/53 (47.17) | 7/53 (13.21) |
| VDE 3y | 97 | 48/97 (49.48)<sup>ab</sup> | 12/97 (12.37) |
| VDE 4y | 49 | 26/49 (53.06)<sup>a</sup> | 5/49 (10.20) |
| 100C | 100 | 24/100 (24)<sup>b</sup> | 9/100 (9)<sup>a</sup> |

The same superscript letters in the same column correspond to a significant difference between experimental groups at $P<0.05$ . FS, frozen semen; VDE, vacuum-dried encapsulated; VDE 0, VDE spermatozoa at time zero; VDE 2y, VDE spermatozoa stored at room temperature (RT) for two years; VDE 3y, VDE spermatozoa stored at RT for three years; VDE 4y, VDE spermatozoa stored at RT for four years; 100C, VDE spermatozoa stored at RT for four years and then incubated at 100°C for ten minutes.

The blastocyst rate for VDE spermatozoa stored for 0, 2, 3, and 4 years decreased slightly over time (Pearson index=-0.8, *P*>0.05). Remarkably, oocytes fertilized with spermatozoa stored for four years or heat-treated (100°C for ten minutes) developed to the blastocyst stage at the same rate (9% for heat-treated spermatozoa vs. 10.2% for the 4-year VDE group, *P*>0.05).

### Embryo Quality

The relative gene expression, based on RT-qPCR on three blastocysts per experimental group, showed that while the expression of *Oct4* in FS and 4y was similar (0.97 fold change over FS, *P*>0.05), *Cdx2* and *Inf-τ* in the 4y group were downregulated compared to the FS group (0.25 and 0.11 fold change over FS, respectively, *P*<0.0001; Figure 2g).

**Figure 2.**
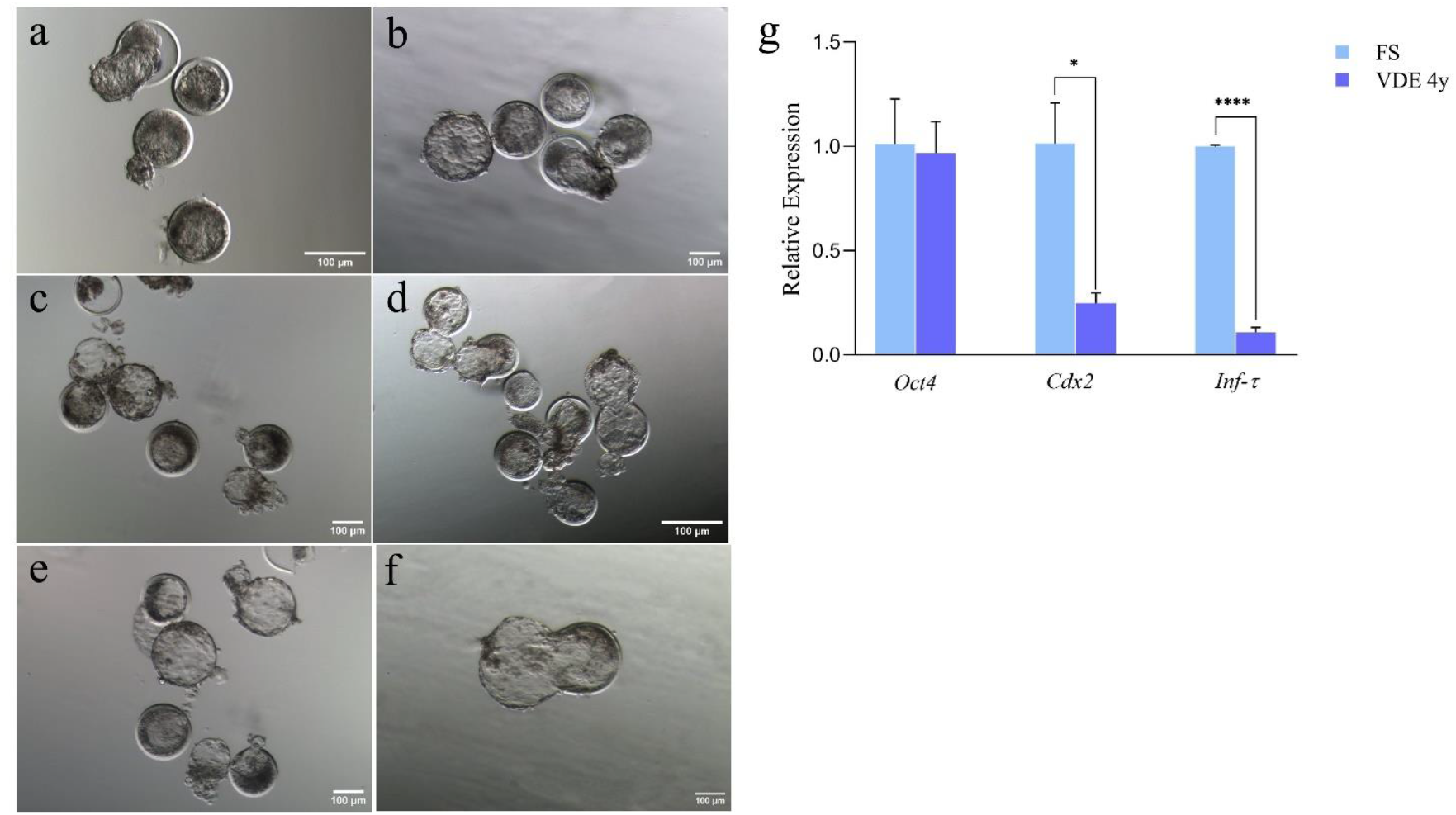
Embryo Quality and Development. **a-f**. Images of day 7–8 embryos. **a**. Following intracytoplasmic sperm injection (ICSI) with FS; **b**. Following ICSI with VDE 0 spermatozoa; **c**. Following ICSI with VDE 2y spermatozoa; **d**. Following ICSI with VDE 3y spermatozoa; **e**, Following ICSI with VDE 4y spermatozoa; **f**. Following ICSI with 100C spermatozoa; **g**. Relative expression *Oct4, Cdx2*, and *Inf-τ* genes in FS and VDE 4y derived embryos; each column represents the median of three replicates; *, *P*<0.05; **** *P*<0.001). FS, frozen semen; VDE 0, vacuum dry encapsulated (VDE) spermatozoa at time 0; VDE 5m, VDE spermatozoa stored for five months at room temperature (RT); VDE 4y, VDE spermatozoa stored for four years at RT; 100C, VDE spermatozoa stored for four years at RT and then incubated at 100°C for 10 minutes.

### Early Pregnancy Loss After Embryo Transfer: Ultrasonography Findings and Follow-up Examinations

Blastocysts of the VDE 4y group were transferred to surrogate mothers to determine whether embryos generated using VDE sperm stored at RT could develop to term. Six adult Sarda ewes (*Ovis aries*) were divided into two groups (three ewes/group). Hatched blastocysts were chosen for embryo transfer on days 7–8 after ICSI.

Two embryo transfer sessions were conducted in each recipient group. Four to five hatched blastocysts were transferred into each recipient sheep, totaling 23 transferred blastocysts. The ultrasonographic examination performed 35–40 days after embryo transfer detected unviable conceptuses compatible with embryo implantation in two recipients. Follow-up ultrasonographic examinations performed a week later confirmed fetal loss.

## DISCUSSION

In this work, we investigated the fertilizing capacity of ram VDE spermatozoa stored at RT for four years. As expected, and similar to other drying methods, the technique compromised sperm vitality and acrosome integrity (Figure 1b-d). Surprisingly, these structural damages were rather stable despite the length of storage (Figure 1d).

Unlike acrosome integrity, DNA damage at the time of dehydration was virtually absent and comparable to the control sample (FS), suggesting that the VDE process had no negative effect on DNA integrity, at least not to an extent greater than sperm freezing, considered the reference method for spermatozoa preservation (Table 1). However, DNA fragmentation increased proportionally with the RT storage time of up to four years (Table 1, Figure 1e).

The overall ICSI results showed a blastocyst rate of 27.59% in the control group (FS) and an insignificant decline in VDE groups over time, decreasing from 13.16% at time 0 to 10.2% after four years of RT storage (Table 2).

VDE 4y spermatozoa were exposed to 100°C for 10 min to realistically estimate the duration they could maintain fertilization capacity when stored at RT [12]. Thermal treatment has been previously utilized to assess the longevity of dehydrated spermatozoa stored at RT [12]. Despite the demonstrated resilience of spermatozoa DNA to high temperatures, they do show a reduced ability to form blastocysts [14]. Heat-stressed spermatozoa were analyzed by SCSA for DNA fragmentation, and their fertilization potential was assessed in ICSI experiments. The outcomes revealed a mild negative correlation between DNA damage and blastocyst development (Figure 1f). The DNA fragmentation in heat-treated spermatozoa remarkably increased from 37.64% to 56.91% (Table 1, Figure 1e), while the blastocyst development rate lowered from 10.2% in the 4y group to 9% in the heat-treated one (Table 2).

Based on the DNA fragmentation/time trend generated using the SCSA empirical data over the first four years of RT storage, we estimated that our accelerated aging trials corresponded to 6.7 years of RT storage.

Although the heat-treated spermatozoa presented an extensive DNA fragmentation rate, blastocysts were obtained (9%). This observation supports our previous findings that highlighted the remarkable DNA repair capabilities of sheep oocytes [17].

We assessed the embryo quality markers gene expression, including that of pluripotent core network gene (*Oct4*), trophoblast transcription factor (*Cdx2*), and interferon tau (*Inf-τ*), the latter a critical prognostic indicator of blastocyst implantation. The results showed that although the blastocysts obtained from VDE 4y spermatozoa did not differ in *Oct4* expression from the control embryos, the expression of *Cdx2* and *Int-τ* underwent a significant downregulation in the 4y group (Figure 2g).

Altogether, these findings demonstrated that the VDE spermatozoa maintained fertilization potential after four years of RT storage. However, sperm DNA damage increased significantly over time, leading to a decline in embryo quality that ultimately compromised embryo implantation. Of the 23 transferred blastocysts derived from VDE spermatozoa stored for four years at RT, only two were implanted, and both were lost within first 40 days. The downregulated genes (*Cdx2* and *Inf-τ*), involved in trophectoderm formation, could partially explain the failure of embryo implantation. In support of this hypothesis, it was reported that paternally-expressed genes predominate in the trophoblast compartment, particularly during placenta development [18]. These trophectoderm impairment findings suggest that VDE trophoblast replacement with FS ones could overcome the inefficient implantation of VDE blastocysts.

In conclusion, the water removal procedure damaged the spermatozoa’s membranes but left their DNA unaffected. The dried sperm storage length and condition (RT) used in our study are unprecedented; even more remarkable is the capacity of these spermatozoa to contribute to embryo development after four years of RT storage. Although we deemed our packaging stable as oxygen was replaced by inert gases before sealing, DNA damage steadily accumulated over time. The residual water might have caused single and double DNA breaks. As suggested above, the oocyte’s DNA repairing activity, which we showed to be highly redundant[17], may have fixed the damage. However, being massive, point mutations were very likely inserted into the genome, exerting their harm during the preimplantation period. This study explored the developmental potential of dry spermatozoa in a farm animal. We believe that our findings encourage further research on dried spermatozoa as a promising preservation method to tackle the decline in biodiversity and simplify male gamete storage, even for ART in humans.

## MATERIALS AND METHODS

### Animals and Ethics Approval

All animal experiments have been approved by the Italian Ministry of Health (no. 200/2017-PR; Prot. 944F0.1 del 04/11/2016). All methods were performed following the relevant guidelines and regulations of the Italian Minister of Health.

### Chemicals

Unless otherwise stated, all materials were purchased from Sigma Aldrich (St Louis, MO, USA)

### Oocyte Collection and *In* Vitro Maturation (IVM)

Sheep ovaries were collected from local slaughterhouses and transported immediately at 37°C to our laboratory, where they were isolated and placed in phosphate-buffered saline (PBS) solution. Cumulus-oocyte complexes (COCs) were aspirated using a 21-gauge needle attached to a 5 mL disposable syringe in the presence of 4-(2-hydroxyethyl)-1-piperazineethanesulfonic acid (HEPES)-buffered TCM-199 medium (Gibco, Life Technologies, Milan, Italy) and 0.005% (w:v) heparin. Only COCs with at least 2-3 layers of clear and compact cumulus cells and uniform ooplasm were selected for IVM. IVM was performed in 4-well dishes containing 500 µL of IVM medium composed of bicarbonate-buffered TCM-199 (Gibco) containing 2 mM glutamine, 0.3 mM sodium pyruvate, 100 μM cysteamine, 10% fetal bovine serum (FBS; Gibco), 5μg/mL follicle-stimulating hormone (FSH) (Ovagen, ICP, Auckland, New Zealand), 5 μg/mL luteinizing hormone (LH), and 1 μg/mL β-estradiol. *In vitro* maturation was completed after incubation for 24 hours in a humidified atmosphere at 38.5°C with 5% CO_2_ in air [19]. Only metaphase II (MII) oocytes with expanded cumulus cells, normal morphology, and polar body I extrusion were selected for ICSI. Cumulus cells were removed by fast pipetting of the COCs in 500 µL of 300 U/mL hyaluronidase solution in HEPES-buffered TCM-199 + 0.4% BSA (w:v). Subsequently, the oocytes were washed three times in HEPES-buffered TCM-199 + 0.4% BSA (w:v). and incubated in a Petri dish pending injection.

### Semen Collection

Semen was collected from an adult fertile Sarda ram using an artificial vagina (AV) filled with warm water (40–44 °C) and connected to a 15-mL collection tube. Immediately after collection, sperm motility was evaluated under a stereomicroscope and transported to our laboratory in a transportable incubator (INC-RB1 Biotherm, Cryologic, Blackburn, Australia) at 38°C.

### Semen Cryopreservation

Semen cryopreservation was performed as previously described [20]. First, a basic medium comprising 2.42 g TRIS base, 1.36 g citric acid, 1.00 g fructose, 100.000 IU penicillin G, and 0.1 g streptomycin in 67.20 mL bi-distilled water (ddH2O); was prepared and adjusted to pH 6.7–6.8. The basic medium was divided into two equal volumes: Medium A, to which 20% egg yolk and 12.8% ddH2O were added, and Medium B, to which 20% egg yolk and 12.8% glycerol were added. An equal volume of each of the two mediums was added to the ejaculates to reach a final concentration of 400 x 10^6^ spermatozoa/mL. The ejaculates, suspended in Medium A (30°C), were incubated at 4°C for two hours. Subsequently, Medium B (4°C) was added, and the suspension was incubated for another two hours at 4°C. The tubes were agitated every 30 minutes. Finally, 250-µL straws were filled, frozen in liquid nitrogen vapors for 20 minutes, and plugged into liquid nitrogen for storage.

### Sperm Vacuum Drying, Encapsulation, and Storage

The spermatozoa, suspended in the Basic Medium, were transferred at 4°C to Imagene Company (France). Upon arrival, the tubes were centrifuged, and the pellet was resuspended in the desiccation medium. Aliquots (50 µL) were placed in stainless steel minicapsules and dried under vacuum for 55 minutes in an evapo-concentrator (HT4, GeneVac, Ipswich, UK). The minicapsules were then transferred into a glove box and maintained for 72 hours under an anoxic and anhydrous argon/helium atmosphere for further desiccation. At the end of the process, the minicapsules were capped with stainless steel caps and sealed by laser welding. Finally, the minicapsules were checked for leakage by mass spectrometry. The ready minicapsules (VDE) were shipped to Italy (University of Teramo) at RT. Samples were stored in the dark for up to four years at RT, pending their use in the experiments.

### Sperm Accelerated Aging Treatment

Bi-distilled water was warmed to 100°C on a heating plate (Tehtnica ROTAMIX 560 MMH). Water temperature was controlled using a Thermometer (ebro handheld TFN 520 TC Type K). VDE microcapsules stored for four years at RT (VDE 4y) were plunged into the 100°C water (100C) and incubated for ten minutes just before being used for ICSI.

### Thawing and Preparation of Frozen Ram Semen

Frozen ram semen (FS) was fast thawed by immersing the straw in 38°C water for 20 seconds. The straw contents were directly transferred into a bicarbonate-buffered synthetic oviductal fluid (SOF) containing 0.4% BSA and spun for 5 min at 120 *g*. The supernatant was removed, and the pellet was resuspended in 100 µL of H199 medium.

### Opening the Minicapsules and Rehydration of the VDE Spermatozoa

VDE minicapsules, stored for 0 and 5 months, and 2, 3, and 4 years (VDE 0, VDE 5m, VDE 2y, VDE 3y, and VDE 4y, respectively) and 100C were opened using an ad hoc shellOpener. The minicapsules were placed in the special housing of the shellOpener, and the shellOpener lid was screwed, ensuring that the tip on the inside of the lid punched a hole in the minicapsules (Figure 1a). All samples were rehydrated with 50 µL bi-distilled water.

### Semen Quality Evaluation

We analyzed membrane and acrosome integrity and DNA fragmentation to determine whether VDE was a valid desiccation technique for storing sperm samples at RT. The samples analyzed included VDE 0, VDE 5m, VDE 4y, and 100C.

#### Flow Cytometry Analysis

Flow cytometry analysis followed the recommendations of the International Society for Advancement of Cytometry [21].

Reagents for flow cytometry were obtained from Thermo Fisher Scientific (Waltham, MA, USA). In each assay, concentration was adjusted to 1 × 10^6^ spermatozoa per mL in a final volume of 0.5 mL Tyrode’s medium, and the spermatozoa were stained with the appropriate combinations of fluorochromes (described below).

Once stained, samples were run through a FACSCalibur flow cytometer (Becton Dickinson, Milan, Italy) with a 488 nm argon-ion laser. The fluorochrome emissions were detected using filters: 530/30 band-pass (green/FL1), 585/42 band-pass (orange/FL2), and >670 long-pass (far red/FL3). Data were acquired using BD CellQuest Pro software (Becton Dickinson).

Fluorescent signals were logarithmically amplified, adjusting the photomultiplier settings to each staining method. FL1 was used to detect green fluorescence from fluorescein-labeled *Pisum sativum* agglutinin (FITC PSA) and acridine orange (AO); FL3 was used to detect red fluorescence from propidium iodide (PI) and acridine orange (AO).

Non-DNA containing particles (debris) in the FITC PSA-stained samples were separately detected by SYBR14/PI analysis, and the percentage of unstained events (PI^−^/FITC PSA^−^) was mathematically determined as described by Petrunkina et al. [22]. Side (SSC-H) and forward (FSC-H) scatter heights were recorded in linear mode (in FSC vs. SSC dot plots). The sperm population was positively gated based on the FSC and SSC, while other events were gated out. At least 10,000 gated sperm events were evaluated per replicate.

#### Membrane and Acrosome Integrity

Sperm acrosome integrity assay (FITC PSA/PI) was used to evaluate membrane and acrosome integrity. Samples were stained with 2.5 µL of PI (Invitrogen Molecular Probes, Eugene, OR, USA; 2.4 mM working solution) and 0.05 mg/mL FITC PSA, incubated at 37°C in the dark for 10 min, and then analyzed.

Four sperm subpopulations were detected on the FL1/FL3 dot plot based on their intact or damaged cellular membrane (negative or positive PI staining, respectively) and intact or damaged acrosome (negative or positive FITC-PSA staining, respectively).

#### Live-dead Staining

As a routine activity in our laboratory, we performed a colorimetric analysis under a fluorescence microscope before processing the samples for flow cytometry. After rehydrating the VDE spermatozoa with 50 µL of bi-distilled water, samples were incubated with 5 μg/mL PI (to detect spermatozoa with a damaged cellular membrane) in PBS for 5 min at RT. Subsequently, a 10-μL aliquot was placed on the slide, covered with a coverslip, and observed on an epifluorescence microscope (Eclipse E-600, Nikon, Tokyo, Japan).

#### PSA Evaluation for Acrosome Integrity

As a routine activity in our laboratory, we performed a colorimetric analysis under a fluorescence microscope before processing the samples in flow cytometry. After rehydrating the VDE spermatozoa with 50 µL bi-distilled water, a smear was made with 10 µL. The slides were air-dried, fixed in 100% ethanol for 5 min, washed three times in PBS, and stained with 200 µL of 1:1 PSA:DAPI (PSA 40 mg/mL, DAPI 0.5 µg/mL, both in PBS). The samples were incubated in the dark for 12 min at RT. Subsequently, the slides were washed 2–3 times in bi-distilled water, mounted with Fluoromount aqueous mounting medium, and analyzed under an epifluorescence microscope (Eclipse E-600, Nikon, Tokyo, Japan).

#### Sperm Chromatin Structure Assay

Sample preparation, processing, and flow cytometer settings were as previously described [23,24]. Briefly, sperm samples were diluted to 200 µL in a buffer solution (0.186 g disodium EDTA, 0.790 g Tris-HCl, and 4.380 g NaCl in 500 mL deionized water, pH 7.4). This was mixed with 400 µL acid detergent solution (2.19 g NaCl, 1.0 mL of 2N HCl solution, and 0.25 mL Triton-X in deionized water to a final volume of 250 mL). After 30 s, 1.2 mL acridine orange solution [3.8869 g citric acid monohydrate, 8.9428 g Na_2_HPO_4_, 4.3850 g NaOH, 0.1700 g disodium EDTA, 4 mg/mL acridine orange stock solution (1 mg/mL) in water to a final volume of 500 mL, pH 6.0] was added. The sample was covered with aluminum foil, placed in the flow cytometer, and allowed to pass through the tubing for 30 s before counting the cells. A total of 5,000 events were evaluated for each sample. Spermatozoa from a single control ram were used as a biological control to standardize the instrument settings between days of analysis. The flow cytometer was adjusted so that the mean green fluorescence was at 500 channels (FL1 at 500) and the mean red fluorescence was at 150 channels (FL3 at 150).

Data were analyzed offline using Kaluza 2.2.1 software. The artificial ‘Sperm DNA Fragmentation’ parameter was calculated using the formula: Red/(Red+Green) fluorescence. The DNA Fragmentation Index (DFI) is expressed as a percentage of spermatozoa with shifted values of the ‘Sperm DNA Fragmentation’ parameter.

### Intracytoplasmic Sperm Injection

VDE 0, VDE 2y, VDE 3y, VDE 4y, and 100C samples were used for ICSI as described before[25]. Briefly, ICSI was performed on an inverted microscope (Eclipse Ti2-U, Nikon) connected to a micromanipulation system (NT-88NEN, Narishige, Tokyo, Japan) and a piezo-driven micropipette system (PiezoXpert, Eppendorf, Milan, Italy). FS and rehydrated VDE spermatozoa (5-μL aliquots) were suspended in 100 μL of H199 with 0.4% BSA (w:v) and then diluted 1:1 with 12% (w:v) polyvinylpyrrolidone in PBS. Three 10–µL drops were placed on the lid of a Petri dish on a warm microscope stage (38.5ºC) and covered with warm mineral oil (38.5ºC). The sperm-containing polyvinylpyrrolidone drops were renewed every ten oocyte injections. After injection, the oocytes were chemically activated by incubation in 5 μM ionomycin in H199 + 0.4% BSA for 5 min, washed once in H199 + 0.4% BSA for 5 min, and then placed in IVC-Medium (BO-IVC, cat. 71005; IVF Bioscience, Falmouth, UK) + 10 µg/mL cycloheximide and incubated for 3.5 hours in humidified atmosphere at 38.5°C and 5% CO_2_. Subsequently, the injected oocytes were washed in IVC-Medium and cultured, as described below.

#### *In Vitro* Embryo Culture

The embryo culture followed our standard laboratory procedure[25] with slight changes. Briefly, fertilized oocytes were cultured in groups of five in 20-µL IVC-Medium (BO-IVC, cat. 71005; IVF Bioscience) drops, covered with mineral oil (Mineral Oil, cat. 51002; IVF Bioscience), and placed in a humidified atmosphere at 38.5°C with 5% CO_2_ and 7% O_2_ for 7–8 days. The *in vitro* development was evaluated for cleavage 24 hours post-activation and on days 7–8 for blastocyst formation. Embryo observation and image acquisition were made on an inverted microscope (Eclipse Ti2-U; Nikon) using Octax EyeWare Imaging Software (version 2.3.0.372; Vitrolife, Västra Frölunda, Sweden).

### Relative Gene Expression

#### RNA Extraction

Three day 7–8 blastocysts from the FS and VDE 4y groups were collected, snap-frozen, and stored in LN pending RNA extraction. Total RNA was extracted using the PicoPure RNA Isolation Kit (Thermo Fisher Scientific, KIT0204) following the manufacturer’s instructions. The PicoPure kit is known for its high efficiency in isolating small amounts of RNA from limited sample material.

#### Complementary DNA (cDNA) Synthesis and Real-time qPCR

The extracted RNA was reverse transcribed into cDNA using the GoScript Reverse Transcription System (Promega A5001) following the manufacturer’s protocol. The obtained cDNA was used as a template to evaluate the relative expression of the pluripotency genes *Oct4, Cdx2*, and *Inf-τ* through real-time PCR amplification. The qPCR was performed in triplicates on the Biorad CFX Connect Real-Time PCR Detection System using SsoAdvanced Universal SYBR Green Supermix (Biorad) following the manufacturer’s instructions. The following gene-specific primers were designed using appropriate bioinformatics tools and following established guidelines: *Oct4*: (*Bos taurus*, NM_174580) forward: aagctcctaaagcagaagagg; reverse: ttctcgttgttgtcagcttcc; *Cdx2*: *Bos taurus*, XM_871005.3) forward: aagacaaataccgggtcgtg; reverse: ctctgcggttctgaaaccaa; *Inf-τ*: *Ovis aries*, XM_027963945.1) forward: ctctgcactggactccaaca; reverse: cgctgtatcccttctcttgc. The thermal cycling conditions were 95°C for 30 s, followed by 45 cycles of 95°C for 15 s and 60°C for 60 s. Melting-curve analysis was performed at the end of each run using the following thermal conditions: 65°C to 95° at 0.5°C increments every 0.05 s to rule out false-positive signals.

#### Quantification and Data Analysis

Gene expression quantification was done using the comparative threshold cycle (ΔΔCt) method. The housekeeping gene β-Actin was used as an internal control for normalization. The results are presented as fold changes relative to a calibrator sample (or a control group) to account for any potential experimental variations.

### Embryo Transfer

Sarda ewes (*Ovis aries*) without lameness and with adequate body condition score, correct mammary gland development, and good general health were selected. Six adult females were selected and divided into two groups of three.

#### Estrus Synchronization

Animals were synchronized using two 20 mg CRONO-GEST sponges inserted into the vagina for 14 days.

#### Embryo Selection and Preparation

Day 7–8 hatched blastocysts were selected for embryo transfer. Assisted hatching of expanded blastocysts was performed using a laser Octax NaviLaser System (Vitrolife, Västra Frölunda, Sweden, ref. 19310/0146) mounted on an inverted microscope (Eclipse Ti2-U, Nikon).

#### Embryo Transfer

Two embryo transfer sessions were conducted in each sheep group, transferring 4–5 hatched blastocysts per recipient sheep totaling 23 transferred blastocysts. Embryos were loaded into a transfer catheter (catheter et tomcat ovini-caprini (19001/0050, Minitube, Alcyon) and surgically transferred into the uterus of each recipient sheep under sedation and general anesthesia.

#### Pregnancy Diagnosis and Monitoring

Pregnancy diagnosis was performed approximately 35–40 days after each embryo transfer session using an ultrasonographic examination. Pregnant ewes were closely monitored for the successful development of the transferred embryos.

### Statistical Analysis

Data were analyzed using GraphPad Prism for Windows (Version 8.0.1, GraphPad Software, San Diego, CA, USA), and statistical significance was set at *P*<0.05.

The chi-squared test was used to compare groups for embryo development outcomes and PSA data. The Brown-Forsythe and Welch ANOVA tests were applied to analyze the SCSA data.

Welch’s *t*-test was applied to relative gene expression results.

Linear regression analysis was performed on the VDE 0, VDE 5m, and VDE 4y SCSA data to obtain a linear regression curve for spermatozoa DNA fragmentation over time.

A correlation analysis (Pearson index) was performed associating blastocyst rate (%) and DFI% in the VDE 0, VDE 4y, and 100C groups.

Pearson analysis was used to correlate the acrosome damage and blastocyst rate over time.

## Acknowledgments

This work received funding from the European Union’s Horizon 2021–2027 Research and Innovation Program under the Marie Skłodowska-Curie action, project “WhyNotDry,” GA-101131087 by the European Union – Next Generation EU, project code: ECS00000041; VITALITY and MUR Prin 2022CSHPAS and Prin P2022FA79R to PL. MCz and LP received funding from the National Science Centre, Poland, through grant 2019/35/B/NZ3/02856 (OPUS). This publication was carried out in the framework of a doctorate co-financed by the European Union – FSE REACT-EU, PON Research and Innovation 2014–2020.

## CRediT Authorship Contribution Statement

**Margherita Moncada:** Writing – Original Draft, Writing – Review & Editing, Conceptualization, Methodology, Validation, Formal analysis, Investigation, Visualization. **Martina Lo Sterzo:** Writing – Original Draft, Writing – Review & Editing, Conceptualization, Methodology, Validation, Formal analysis, Investigation, Visualization. **Luca Palazzese:** Writing – Review & Editing, Conceptualization, Investigation, Visualization, Supervision. **Francesca Boffa:** Investigation. **Debora Agata Anzalone:** Investigation. **Domenico Iuso:** Writing – Review & Editing, Visualization, Supervision. **Marta Czernik:** Writing – Review & Editing, Resources, Project administration, Funding acquisition. **Luisa Gioia:** Writing – Review & Editing, Resources. **Federica Turri:** Investigation. **Beatrice Mislei:** Writing – Original Draft, Investigation. **Diego Bucci:** Investigation. **Jacques Bonnet:** Investigation. **Marthe Colotte:** Investigation. **Sophie Tuffet:** Investigation. **Michał Bochenek:** Investigation. **Pasqualino Loi:** Writing – Review & Editing, Resources, Project administration, Funding acquisition.

## Data Availability Statement

Data will be made available on request.

